# Aberrant expression of CD14 in breast tumor cells is associated with poor outcome

**DOI:** 10.1101/2024.11.07.622456

**Authors:** Hiroyuki Katayama, Rongzhang Dou, Ricardo A. León-Letelier, Ehsan Irajizad, Alejandro Sevillano, Soyoung Park, Fu Chung Hsiao, Yining Cai, Jody Vykoukal, Johannes Fahrmann, Jennifer Dennison, Elizve Barrientos-Toro, Maria Gabriela Raso, Aysegul Sahin, Sam Hanash

## Abstract

Emerging evidence suggests that cancer cells can mimic features of immune cells during oncogenic transformation to drive disease progression. We assessed the occurrence of immunological markers in breast cancer cells to determine their expression pattern. We initially analyzed 18 immune protein markers (CCR4, CCR6, CCR7, CD11, CD123, CD14, CD16, CD19, CD24, CD25, CD27, CD3, CD38, CD4, CD45, CD56, CD8 and CXCR3) expressed on the surface of 28 breast cancer cell lines using mass spectrometry. CD14 protein expression in tumor cells and its association with clinical outcomes was subsequently evaluated by tissue microarray (TMA) analysis of 346 breast tumors. Single-cell RNA sequencing data from breast cancer tumors and bulk transcriptomic data of breast cancer cell lines were interrogated for molecular signatures associated with CD14 tumor cell expression. Among the markers interrogated, CD14 *protein* was aberrantly expressed on the surface of 13 out of 15 triple-negative breast cancer (TNBC), one of six hormone receptor positive (HR+), one of five hormone receptors negative (HR-)/Her2+ cell lines. Likewise, RNA expression revealed higher levels of CD14 in TNBC cell lines compared to other subtypes. Tumor tissue microarray analysis revealed elevated levels of CD14 membrane expression predominantly in TNBC and was associated with higher tumor grade and increased incidence of disease recurrence compared to CD14-negative tumors. The CD14-positive subgroup exhibited Nuclear Factor Kappa Beta (NFkB) and TGF-β centric networks at both the protein and Single-cell RNA levels. We have uncovered a novel subset of breast cancers characterized by aberrant surface expression of CD14 associated with aggressive disease. CD14 identifies a subset of breast cancers with poor outcome and is a potential therapeutic target.

## Introduction

Breast cancer is largely defined based on the expression of estrogen receptor (ER), progesterone receptor (PR) and epidermal growth factor receptor 2 (Her2). Triple negative breast cancer (TNBC) which consists of 10 to 15 % of breast cancers exhibits an aggressive phenotype with fewer therapeutic options(1). The Her2-enriched subtype is also associated with a poor prognosis(2). There is a need to identify additional surface markers for molecular subtyping beyond classical hormone receptor classification with a potential to refine classification and expand therapeutic options.

We assessed whether the concept of immune mimicry-through expression of immune markers has relevance to breast cancer. A subset of epithelial cells was found to mimic regulatory T cells and contribute to immune evasion in pancreatic adenocarcinoma(3,4). In other studies, the monocyte marker CD14 was found to be expressed in bladder cancer cells establishing an inflammatory and proliferative prone tumor microenvironment(5). Other surface proteins expressed in immune cells were found to be expressed in cancers including the macrophage marker CD36 in acute myeloid leukemia and in colorectal and gastric cancers(6), the macrophage and dendritic cells marker CD58 in melanoma(7), the T cell marker CD70 in non-small cell lung cancer and glioblastoma(8–10), T cell and NK cell marker CD160 in melanoma and esophageal adenocarcinoma(11,12), and the dendritic cell marker CD320 in gastric cancer(13).

We analyzed in this study expression of immune surface markers initially in breast cancer cell lines using proteomics which led to the discovery of the monocyte marker CD14 expression. Analysis of CD14 in tissue microarrays (TMA) was undertaken to determine its expression across subtypes and its relevance to outcome. CD14-positive (CD14 (+)) and CD14-negative (CD14 (−)) subgroups were compared in available, Single-cell RNA sequencing (RNA-seq) data for related networks.

## Methods

### Proteomic analysis

A total of 26 breast cancer cell lines MCF7 (Cat# HTB-22, RRID: CVCL_0031), HCC1500 (Cat# CRL-2329, RRID: CVCL_1254), T47D (Cat# HTB-133, RRID: CVCL_0553), ZR75-1 (Cat# CRL-1500, RRID: CVCL_0588), CAMA1 (Cat# HTB-21, RRID: CVCL_1115), BT474 (Cat# HTB-20, RRID: CVCL_0179), HCC1954 (Cat# CRL-2338, RRID: CVCL_1259), HCC202 (Cat# CRL-2316, RRID: CVCL_2062), HCC2218 (Cat# CRL-2343, RRID: CVCL_1263), MDA-MB-361 (Cat# HTB-27, RRID: CVCL_0620), SKBR3 (Cat# HTB-30, RRID: CVCL_0033), MDA-MB-231 (Cat# HTB-26, RRID: CVCL_0062), HCC1143 (Cat# CRL-2321, RRID: CVCL_1245), HCC1937 (Cat# CRL-2336, RRID: CVCL_0290), HCC1599 (Cat# CRL-2331, RRID: CVCL_1256), HCC1806 (Cat# CRL-2335, RRID: CVCL_1258), MDA-MB-468 (Cat# HTB-132, RRID: CVCL_0419), HCC70 (Cat# CRL-2315, RRID: CVCL_1270), HCC1187 (Cat# CRL-2322, RRID: CVCL_1247), HS578T (Cat# HTB-126, RRID: CVCL_0332), BT549 (Cat# HTB-122, RRID: CVCL_1092), HCC1395 (Cat# CRL-2324, RRID: CVCL_1249), HCC38 (Cat# CRL-2314, RRID: CVCL_1267), MDA-MB-436 (Cat# HTB-130, RRID: CVCL_0623), BT20 (Cat# HTB-19, RRID: CVCL_0178), MDA-MB-157 (Cat# HTB-24, RRID: CVCL_0618) were obtained from ATCC (American Type Culture Collection, Manassas, VA, USA) and HMLER2, HMLER3, BPLER2 and BPLER3 were obtained from Dr. Susan Lindquist’s group. All ATCC cell lines were authenticated using short tandem repeat (STR) profiling by ATCC. The HMLER and BPLER cell lines were authenticated by Dr Susan Lindquist’s group. All cell lines were tested for mycoplasma contamination using PCR-based assays before experimentation. Trypsin digestion was applied to surface proteins and total cell extracts (TCE) followed by nano-liquid chromatography (nanoLC)-mass spectrometry as previously described(14–19).

### RNA-seq analysis

RNA-seq gene expression data of breast cancer cell lines was obtained from CCLE-Cancer Cell Line Encyclopedia (www.sites.broadinstitute.org/ccle/). Gene expression transcripts per million (TPM) values were inferred from RNA-seq data using RSEM software package(20) and reported after log2 transformation, using pseudo-count of 1; log2(TPM+1). A total of 66 cell lines were classified as, hormone receptor positive (HR+) subtype (21 cell lines) consisting from MDA-MB-415, BCK4, MCF7, UACC812, EFM19, MDA-MB-157VII, CAMA1, MDA-MB-361, HCC1500, EMF192A, KPL1, SUM44PE, MDA-MB-134VI, BT483, SUM52PE, HCC1428, T47D, ZR-7530, MFM223, ZR75-1 and BT474, hormone receptor negative and Her2 positive (HR-/Her2+) (13 cell lines) consisting of JIMT1, SUM229PE, HCC1954, COLO824, UACC3133, AU565, UACC893, HCC1569, SKBR3, HCC1419, HCC202, SUM190PT and HCC2218, and TNBC (32 cell lines) consisting of HCC70, HCC1937, MDA-MB-468, MDA-MB-157, HCC1187, HCC2157, HCC1143, MDA-MB-436, VP267, SUM149PT, HCC38, SUM159PT, VP229, HC1599, HCC1806, HS578T, CAL120, CAL851, SNU2372, BT20, MDA-MB-21, DU4475, HCC1395, HDQP1, BT549, CAL51, CAL148, SUM185PE, MDA-MB-453, HMC18, SUM102PT and SUM1315MO2. The reference knowledge was utilized for classification of those hormone receptor subtypes(21,22).

### Tissue microarrays

Tissue microarrays (TMAs) consisting of a total of 346 newly diagnosed HR+, HR-/Her2+ and TNBC tumors were analyzed for CD14 expression (Supplementary Table 7). The study was approved by the IRB at MD Anderson Cancer Center. Tissue slides of normal skin and spleen (US Biomax, Cat# BR1301, Rockville, MD, USA), mammary gland tissues (Zyagen, Cat# HP-414, San Diego, CA, USA) and normal tonsil were obtained from MD Anderson. The TNBC, MDA-MB-468 (Cat# HTB-132, RRID: CVCL_0419) cells were obtained from ATCC and prepared for IHC.

### Immunohistochemistry workflow

Breast cancer cell line and normal tissue slides were de-paraffinized in xylene, rehydrated in a descending ethanol series, and subsequently treated with 3% hydrogen peroxide for 10 min. Antigen retrieval was conducted in a pressure cooker in 20x ImmunoDNA Retriever with citrate (Bio SB, Cat# BSB0022, Santa Barbara, CA, USA) and 0.1 % Tween 20 at 121°C for 15 minutes. Sections were hybridized with 1:3,000-times diluted anti-CD14 monoclonal antibody (clone 1H5D8, Abcam, Cat# ab181470, Camdridge, MA, USA) for 16 hours at 4°C. After washing with TBS for 5 minutes x3, signal development was performed with Histofine DAV-2V kit (Nichirei bioscience, Cat# 425312F, Tokyo, Japan). The breast cancer TMA slides were stained using an automated stainer Leica BOND Max, 4 µm-thick TMA slides were incubated with the primary CD14 monoclonal antibody (clone 1H5D8, Abcam, Cat# ab181470, Camdridge, MA, USA) at a dilution of 1:3,000 for 15 minutes at room temperature. Pretreatment and antigen retrieval were performed at pH 6 for 20 minutes. Images were scanned with MDACC-North campus Research Histology Core Laboratory and analyzed using Aperio Imagescope (Leica Biosystems, Buffalo Grove, IL, USA). For IHC evaluation, p-values for association with CD14 positive versus negative expression were calculated based on 2-sided kaisquare test.

### Single-cell analysis

TNBC single-cell RNA-seq data was retrieved from the Single-Cell Portal website (https://singlecell.broadinstitute.org/single_cell), specifically, from data uploaded by Wu et al(23). The Seurat package (version 4.3.0) (24) implemented in R statistical software (version 4.3) (https://www.r-project.org/) was used for data-filtering, pre-processing and principal component analysis (PCA) based-dimension reduction and t-distributed stochastic neighbor embedding (t-SNE) to separately visualize the cell clusters from tumor microenvironment. The expression and percentage of CD14 was calculated accordingly.

### Network and statistical analyses

Networks analysis was done using Ingenuity Pathway Analysis (www.ingenuity.com/)(25). Figures and Tables were generated using R statistical software or GraphPad Prism software V10 (www.graphpad.com/). Survival analysis (Table 3) was performed in R software (version 3.6.1, The R Foundation, https://www.r-project.org) using survival package. The 95 % confidence interval (C.T.) and corresponding p values were calculated by 10,000 bootstrap samples. A Cox model was applied to construct survival curves and to estimate hazard ratios; the log-rank test was used to evaluate the statistical significance. To test for the proportionality of Hazard assumption of a Cox regression, we utilized the method of Grambsch et al(26). In the Kaplan-Myer plot of Curtis’ gene expression data(27) (Supplementary Fig. S2), CD14^high^ versus CD14^low^ groups were based on median cut-off.

## Data availability

The data generated in this study are available within the article and its supplementary data files.

## Results

### CD14 expression in breast cancer cells

We initially conducted proteome analysis of breast cancer cell line surface proteins and total cell extracts (TCE) using mass spectrometry to assess expression of immune-associated markers. The analysis covered immunophenotype markers from the Human Immunology Project(28) that included 18 immune markers (CCR4, CCR6, CCR7, CD11, CD123, CD14, CD16, CD19, CD24, CD25, CD27, CD3, CD38, CD4, CD45, CD56, CD8 and CXCR3) (Supplementary Table S1). The monocyte marker CD14, was found to be expressed on the surface of one out of six HR+, one out of five HR-/HR2+ and 13 out of 15 TNBC, total 28 cell lines and exhibited surface enrichment compared to TCE (Fig. 1 (A), Fig. 1(B), Supplementary Table S2-S3). Other markers investigated included the NK cell marker CD56 (NCAM1) which was identified on the cell surface of one HR+ cell lines, one HR-/Her2+ cell lines and six TNBC cell lines (Fig. 1 (A), Fig. 1(B), Supplementary Table S2-S3). Given the prominence of CD14 among 18 immune markers, we further investigated RNA-seq transcriptomic data available from CCLE (www.sites.broadinstitute.org/ccle/) for total of 66 breast cancer cell lines. CD14 was expressed at the transcriptomic level in 32 TNBC, 21 HR+ and 13 HR-/Her2+ subtypes (Fig. 1 (C), Supplementary Table S4).

**Fig. 1.**
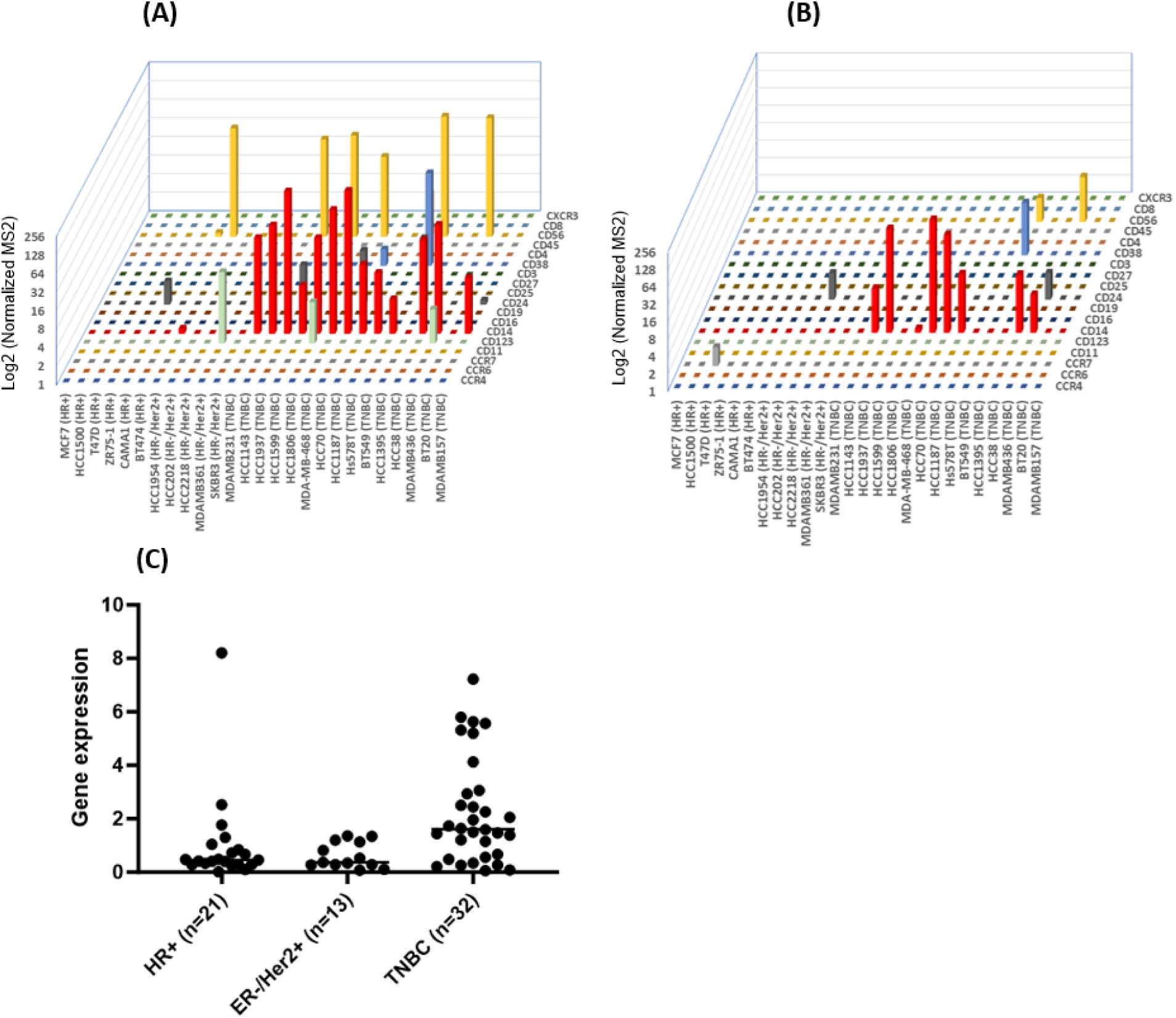
Immune-associated markers expression in breast cancer cell lines. (A) Dominant expression of CD14 protein on cell surface. (B) Dominant expression of CD14 protein in Total Cell Extracts (TCE). (C) CD14 mRNA expression in RNA-seq (CCLE) of (n=66) breast cancer cell lines.

### CD14 expression in breast tumor tissues and clinical outcome

We assessed CD14 expression across hormone receptor subtypes and its correlation with clinical outcome in breast tumor tissues. CD14 staining of the cell line MDA-MB-468 (TNBC) was used as a positive control (Fig. 2(A)) and staining of normal mammary gland, skin, and tonsillar tissues as a negative control (Supplementary Fig. S1).

**Fig. 2.**
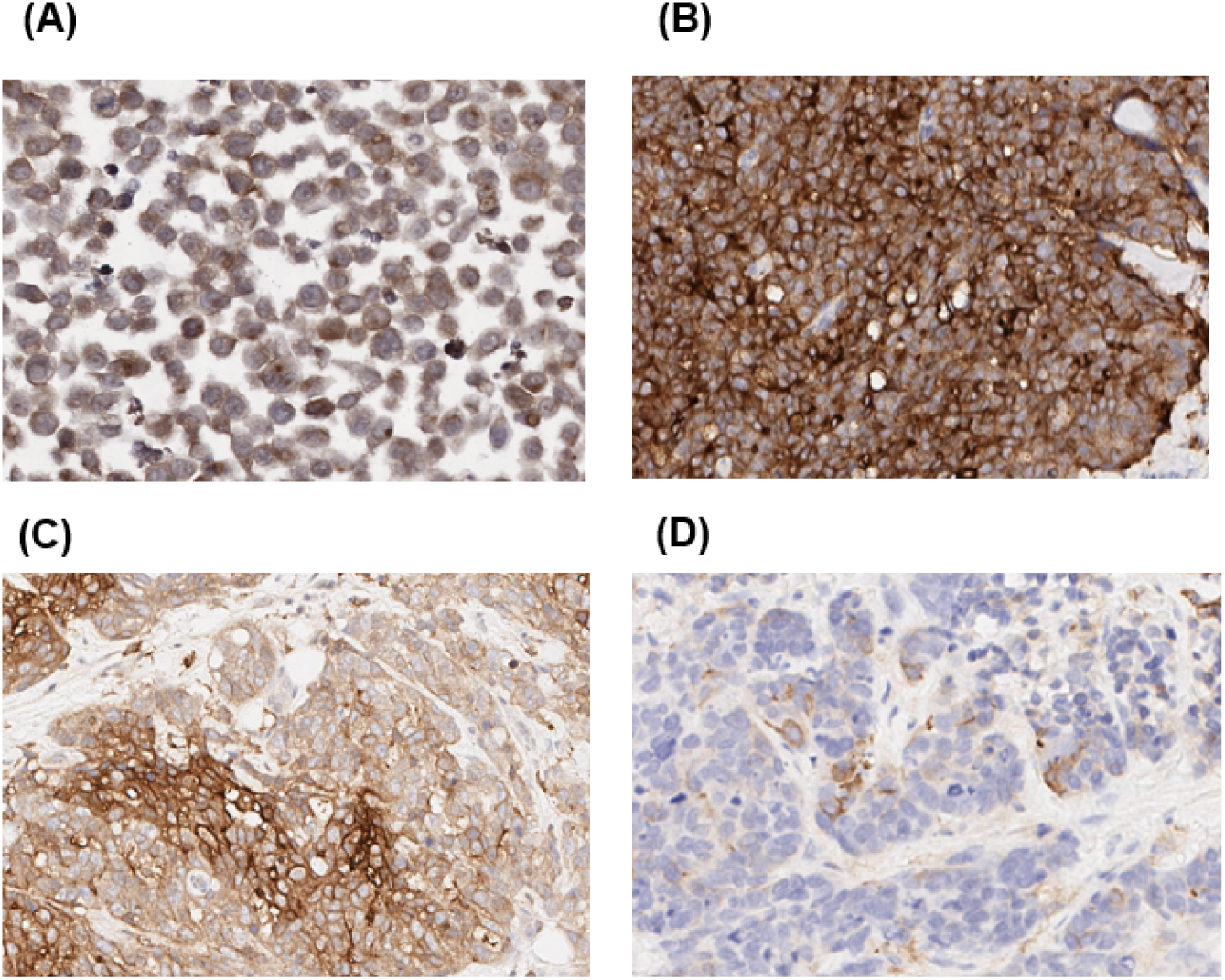
Immunohistochemistry (IHC) of CD14 on breast cancer cell surface. (A) anti-CD14 antibody IHC in MDA-MB-468 (TNBC) cells showing positive surface staining. (B) anti-CD14 antibody IHC in TNBC patient (Case 1), showed strong tumor cell surface staining. (C) anti-CD14 antibody IHC in TNBC patient (Case 2,) showed moderate tumor cell surface staining. (D) anti-CD14 antibody IHC in TNBC patient (Case 3), showed weak tumor cell staining.

We assessed CD14 expression on TMAs consisting of 346 breast tumors (Table 1, Supplementary Table S7, Fig 2 (B), (C), (D)). CD14 surface expression was significantly higher in grade III tumors versus I+II (p<0.0001) and in recurrent versus non-recurrent tumors (p=0.0326) (Table 2). Notably, CD14 expression was also detected in a subset of ER(+) tumors.

**Table 1.**
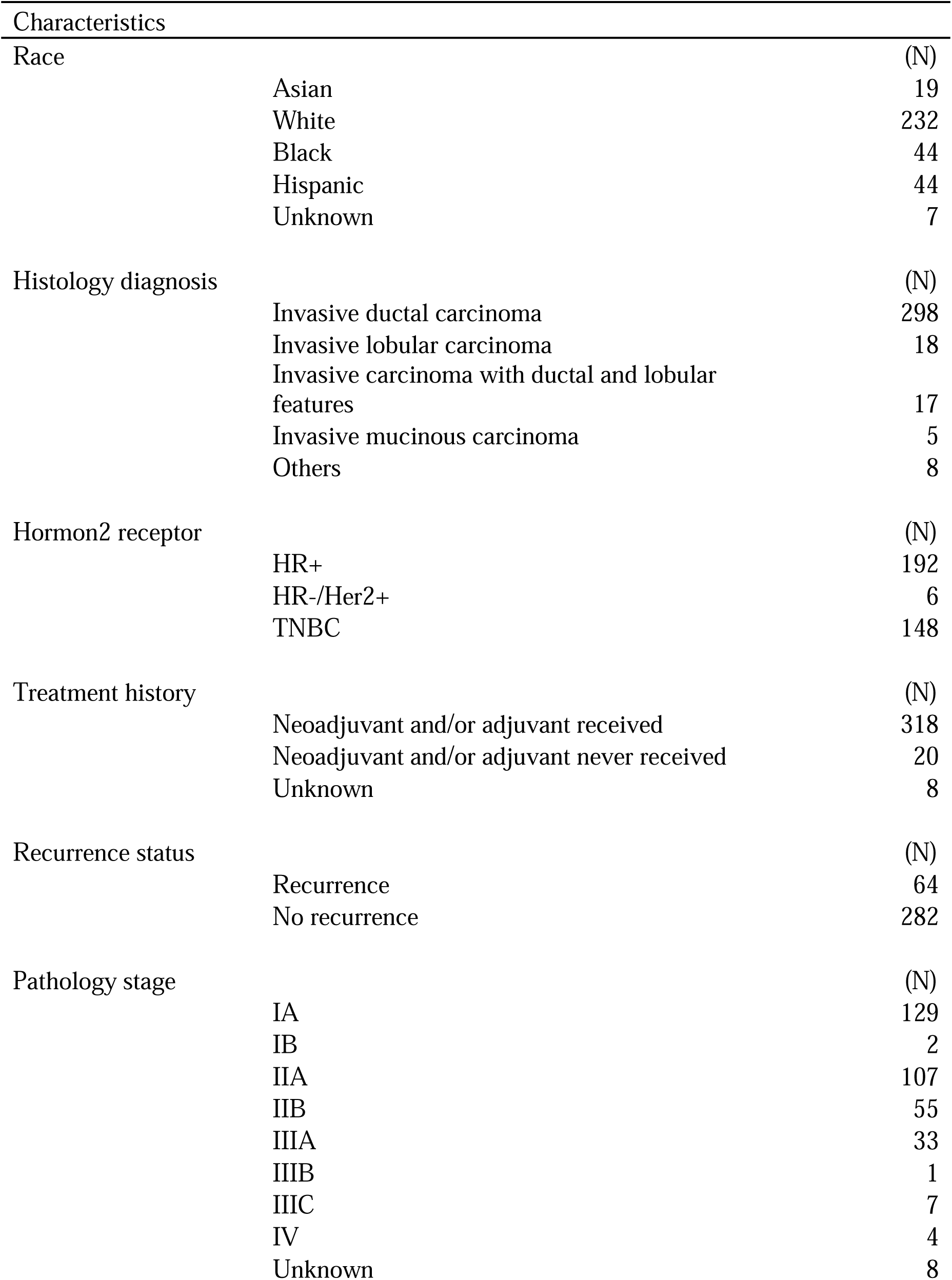
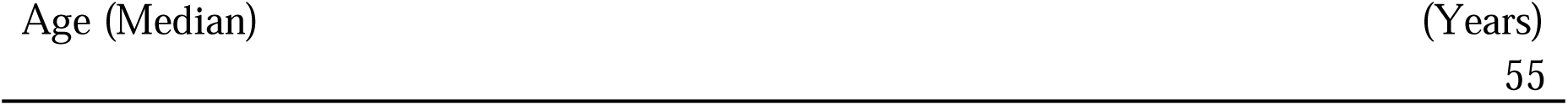
Patient characteristics of breast cancer tissue microarray (N=346)

| Characteristics |  |  |
| --- | --- | --- |
| Race |  | (N) |
|  | Asian | 19 |
|  | White | 232 |
|  | Black | 44 |
|  | Hispanic | 44 |
|  | Unknown | 7 |
| Histology diagnosis |  | (N) |
|  | Invasive ductal carcinoma | 298 |
|  | Invasive lobular carcinoma | 18 |
|  | Invasive carcinoma with ductal and lobular features | 17 |
|  | Invasive mucinous carcinoma | 5 |
|  | Others | 8 |
| Hormon2 receptor |  | (N) |
|  | HR+ | 192 |
|  | HR-/Her2+ | 6 |
|  | TNBC | 148 |
| Treatment history |  | (N) |
|  | Neoadjuvant and/or adjuvant received | 318 |
|  | Neoadjuvant and/or adjuvant never received | 20 |
|  | Unknown | 8 |
| Recurrence status |  | (N) |
|  | Recurrence | 64 |
|  | No recurrence | 282 |
| Pathology stage |  | (N) |
|  | IA | 129 |
|  | IB | 2 |
|  | IIA | 107 |
|  | IIB | 55 |
|  | IIIA | 33 |
|  | IIIB | 1 |
|  | IIIC | 7 |
|  | IV | 4 |
|  | Unknown | 8 |

|  |  |
| --- | --- |
| Age (Median) | (Years) |
|  | 55 |

**Table 2.**
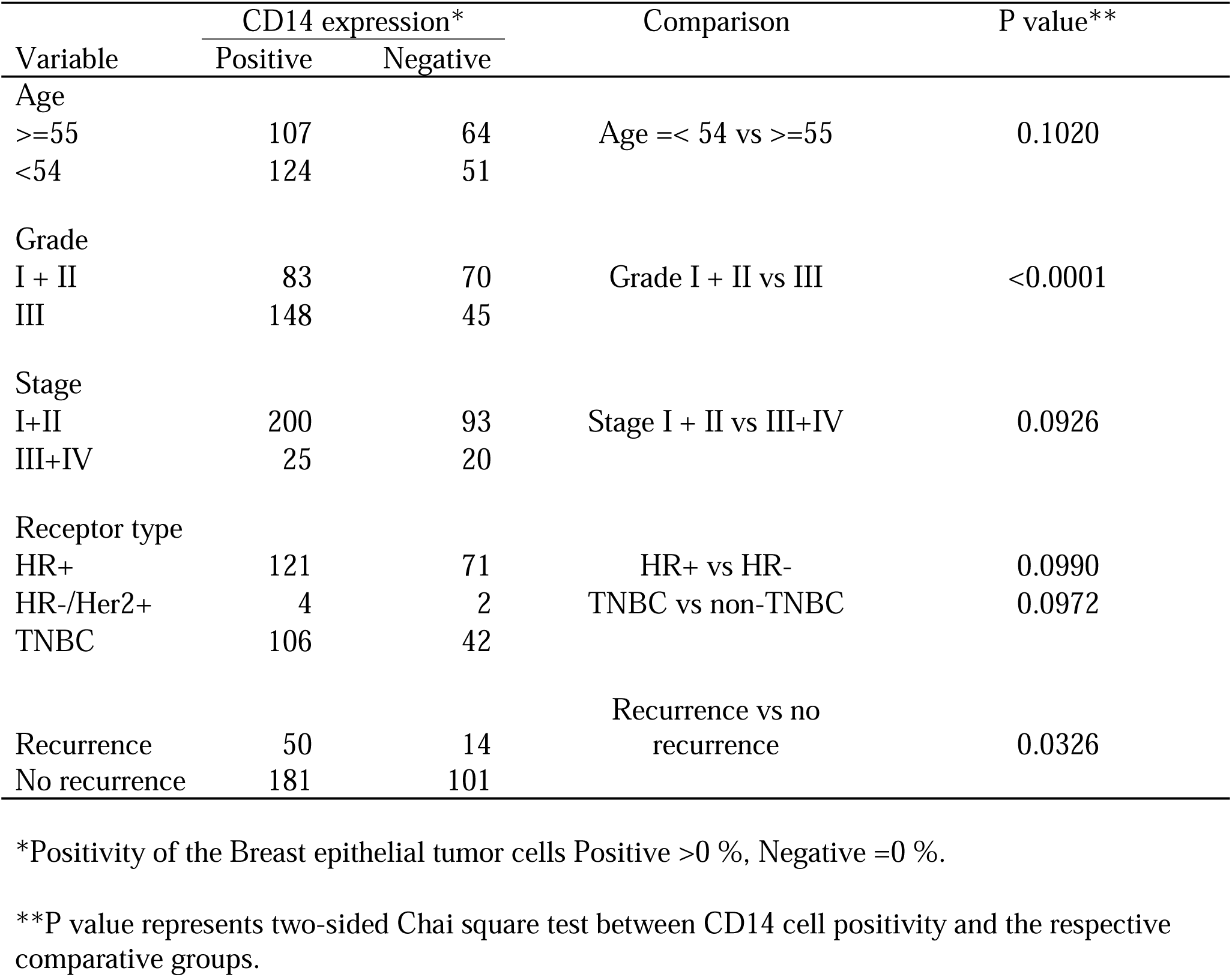
CD14 membrane expression in breast cancer tissues (n=346) in relation to clinical parameters.

Using the optimal cut-point method, we stratified tumors into positive and negative groups. The hazard ratio between the two groups (adjusted for staging) across all breast cancer subtypes was 1.854 (P=0.017). Further stratification based on TNBC and non-TNBC subtypes resulted in stage-adjusted hazard ratio of 1.593 and 2.427, respectively (Table 3).

**Table 3.**
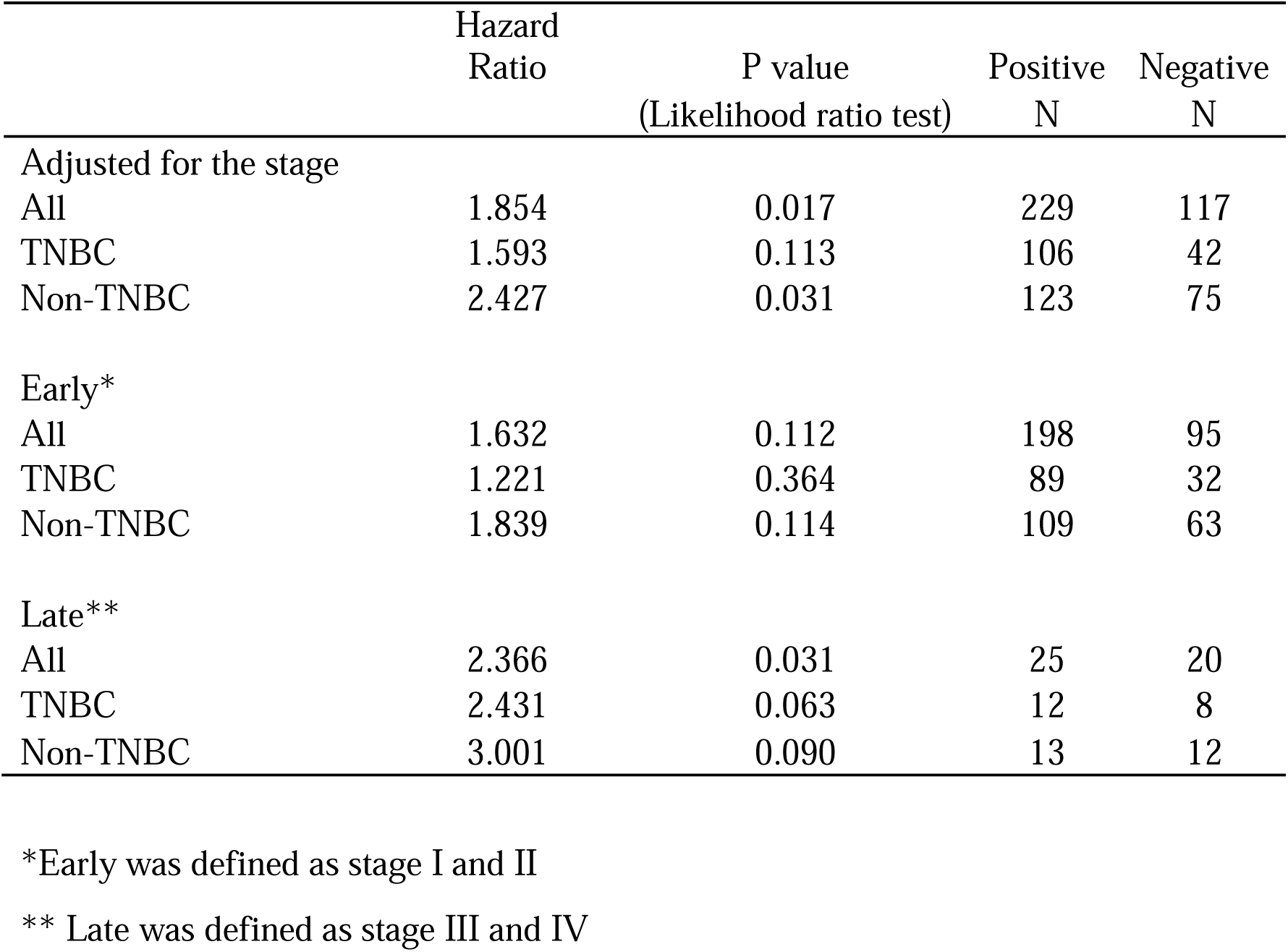
Survival analysis of breast cancer tissue microarray (n=346)

|  | Hazard<br>Ratio | P value<br>(Likelihood ratio test) | Positive<br>N | Negative<br>N |
| --- | --- | --- | --- | --- |
| Adjusted for the stage |  |  |  |  |
| All | 1.854 | 0.017 | 229 | 117 |
| TNBC | 1.593 | 0.113 | 106 | 42 |
| Non-TNBC | 2.427 | 0.031 | 123 | 75 |
| Early* |  |  |  |  |
| All | 1.632 | 0.112 | 198 | 95 |
| TNBC | 1.221 | 0.364 | 89 | 32 |
| Non-TNBC | 1.839 | 0.114 | 109 | 63 |
| Late** |  |  |  |  |
| All | 2.366 | 0.031 | 25 | 20 |
| TNBC | 2.431 | 0.063 | 12 | 8 |
| Non-TNBC | 3.001 | 0.090 | 13 | 12 |

\*Early was defined as stage I and II
\*\* Late was defined as stage III and IV

In the Kaplan Meier plot of Curtis’ gene expression data(27), CD14^high^ was associated with poor outcomes in early stage (stage 0 and stage I) in HR+ (P=0.0327, HR 1.367, 95 % CI 1.024 to 1.825) and TNBC (P=0.0003, HR 5.795, 95 % CI 2.497 to 13.47) (Supplementary Fig. S2 (A)-(C)).

### CD14-associated TGF-**β** signature

Analysis of proteomic TCE data in subgrouping breast cancer cell lines split into CD14 (+) and CD14 (−) revealed networks involving the NFkB complex and TGF-β signaling (P<0.05) (Fig. 3 (A), supplementary Table S5). Analysis of a metastatic and a non-metastatic models, consisting BPLER and HMLER cells derived from the same TNBC tumor revealed elevated levels of heat shock factor 1 (HSF1) in the metastatic BPLERs(29) and TGF-β signature compared to non-metastatic HMLER(15). BPLER cells exhibited higher levels of cell surface CD14 in both BPLER2 vs HMLER2 and BPLER3 vs HMLER3 comparison (Fig. 3 (B)).

**Fig. 3.**
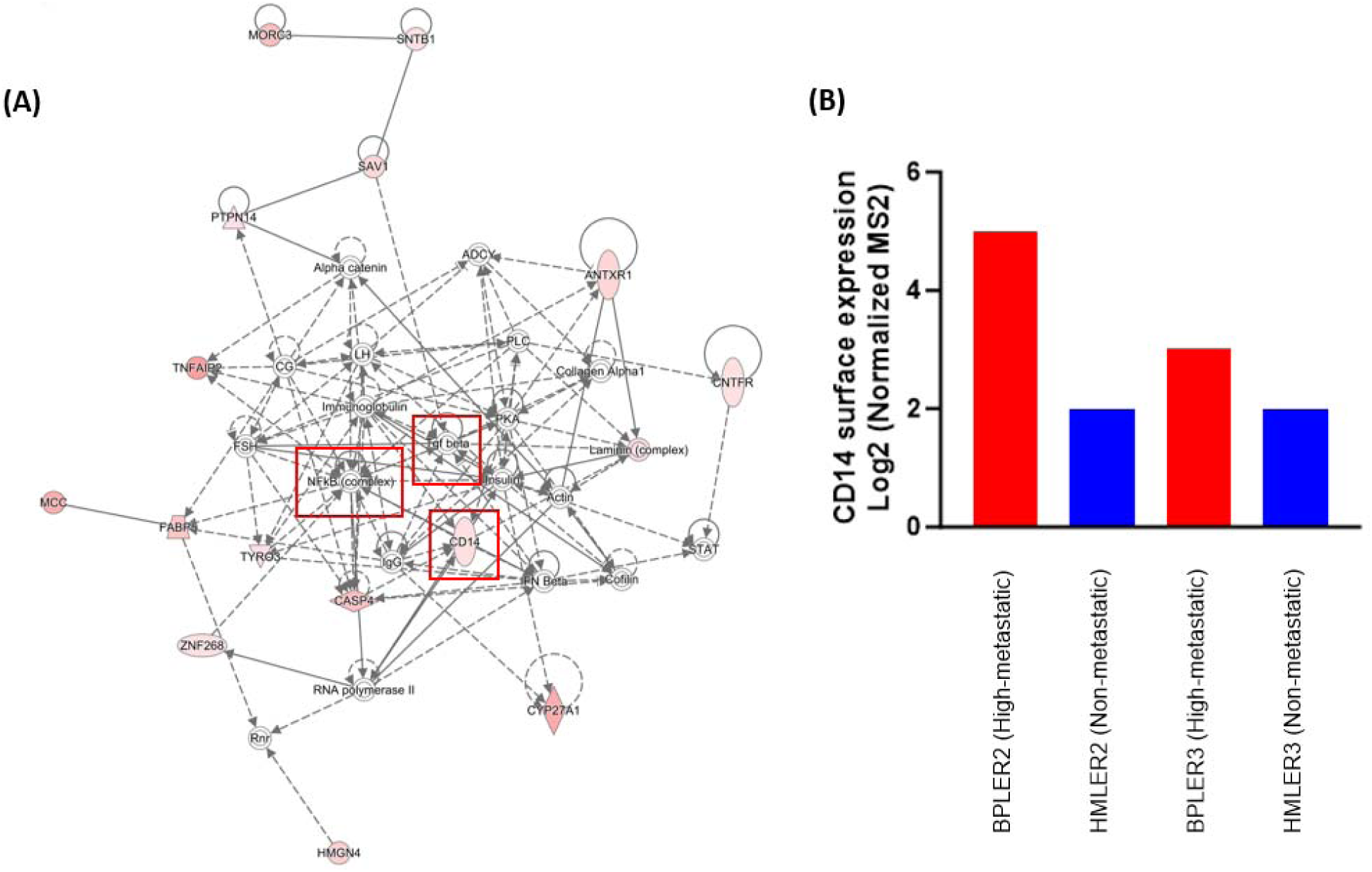
TGF-β signature associated with CD14 in breast cancer. (A) The CD14 (+) compared to CD14 (−) subgroup was associated with TGF-β and NFkB signature in the TCE proteome (>=10 Change Fold; CD14 (+)/CD14 (−), P<0.05,). CD14 (+) was grouped by HCC70, HCC1937, MDA-MB-468, HCC1187, MDA-MB-436, HCC1143, MDA-MB-157, HCC38, MDA-MB-231, HCC1806, Hs578T, HCC1599 and BT549 whereas CD14 (−) was grouped from MCF7, BT20, HCC1954, HCC1395, HCC202, CAMA1, SKBR3, T47D, HCC2218, BT474, ZR75-1, MDA-MB-361 and HCC1500 based on the CD14 surface expression. (B) TGF-β enriched signature in metastatic BPLERs(15) showed higher level of CD14 on cell surface compared to non-metastatic HMLERs.

### CD14 expression at the single cell level

Given the distinct malignant phenotype of TGF-β activated tumors within the estrogen receptor negative subtype (15,30), we analyzed single-cell gene expression data from TNBC tumors available (https://singlecell.broadinstitute.org/single_cell). Analysis, based on data from five patients’ specimens (18,830 single cells), identified CD14 mRNA expression in 7.12 % of epithelial cancer cells (Fig. 4(A)). Utilizing the CellChat package, we delineated cell interaction maps between CD14 (+) and CD14 (−) epithelial cancer cells revealing unique interactions of CD14 (+) cells with macrophages (thick purple lines) while CD14 (−) cells connected with NK cells and T cells (green lines) (Fig. 4 (B)). Gene set enrichment analysis highlighted the involvement of the NFkB signaling pathway (Padj=0.0118) in CD14 (+) cells (Fig. 4 (C)). Ingenuity Pathway Analysis of CD14 correlated proteins in CD14 (+) and CD14 (−) comparison revealed CD14 (+) was associated with NFkB and TGF-β networks (Spearman correlation, >0.25) which was consistent with cell line analysis (Fig. 4(D), Supplementary Table S6).

**Fig. 4.**
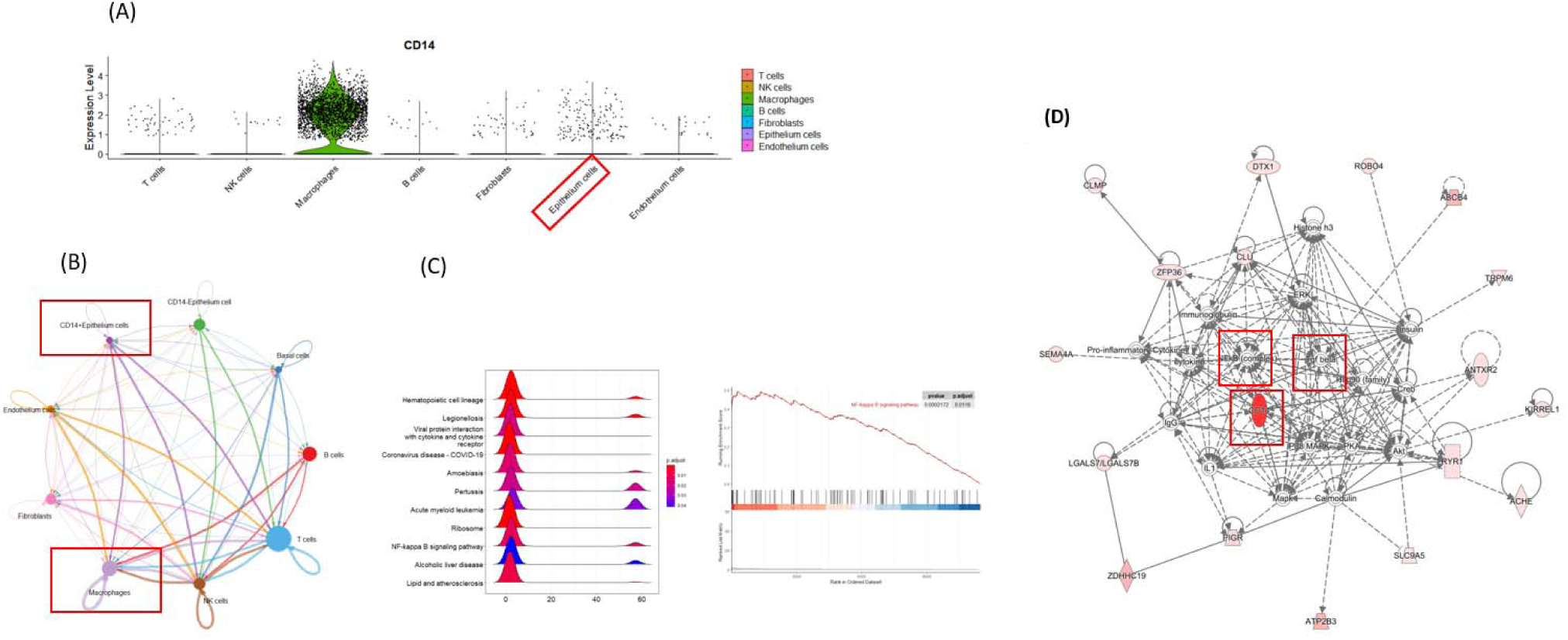
CD14 Single-cell RNA-seq analysis from TNBC patients. (A) Single-cell RNA-seq analysis in TNBC revealed, CD14 was expressed in a subset of the epithelial cancer cells. The cell clusters were defined on the basis of expression of following markers: epithelial cancer cells (KRT19, EPCAM, CDH1, CCND1), endothelial cells (PECAM1, VWF), Fibroblasts (COL1A2, COL1AA, FAP), T cells (CD3D, CD3E, CD2), B cells (CD79A, IGHG1, MS4A1), NK cells (KLRD1, GNLY, KLRF1), macrophages (CSF1R, CSF3R, CD68). (B) Interaction map of CD14 (+) epithelial cancer cells have specific interaction with macrophages (purple lines) compared to CD14 (−) epithelial cancer cells connected to T cells and NK cells (green lines). (C) Pathway enrichment shows significant enrichment of NFkB signaling pathway (Padj=0.0118) in CD14 (+) breast cancer epithelial cancer cells. (D) CD14 (+) associated proteins in breast epithelial cancer cells (Pearson correlation r>0.25) showed NFkB and TGF-β signature.

## Discussion

CD14 is known for its role as a co-receptor for toll-like receptors (TLRs), activating immune responses to pathogens and tissue injury in macrophages and monocytes(31). Our study uncovered CD14 protein expression in breast cancer cells and its association with poor outcome.

CD14 expression in tumor cells has been documented in other cancer types. In cancer cell line studies, CD14 mRNA expression was observed in glioblastoma(32) and gastric cancer(33). In the latter, CD14 knockdown resulted in reduced secretion of TNF-α and TGF-β, and CD14 elevated expression was found to be associated with epithelial to mesenchymal transition (EMT). In bladder cancer, CD90+ basal cells were found to express higher level of CD14(5). Moreover, a significant elevation of inflammatory mediators IL6, IL8 and Macrophage stimulating factor (M-CSF) was observed indicative of upregulation of an inflammatory response. When used in a murine model CD14^high^ cells induced an immune-cold environment with lower degree of MHC II expression on tumor monocytic cells and higher percentage of M2 macrophages in the tumor microenvironment. In the colorectal cancer (CRC) TCGA data, a significant upregulation of CD14 is found in the CMS4 subtype with features of TGF-β activation and EMT. IHC analysis of tumor tissues revealed the CD14^high^ group to have worst outcome (34).

The association between CD14 positivity with an NFkB-TGF-β network both in cell lines and single cell gene expression data derived from TNBC tumors suggests a potential role of CD14 in tumor aggressiveness. TGF-β is a key regulator of inflammation in the tumor microenvironment(35), and plays a crucial role in promoting tumor progression including evasion of immune surveillance, autocrine mitogen and cytokine production, EMT, and myoblast and osteoclast mobilization(36). Molecular profiling of fast and slow-growing tumors from the same strain of ER-breast cancer mouse model revealed TGF-β mRNA level to be significantly higher in the fast-growing tumor group(30). We previously reported, a plasma protein derived TGF-β signature as a prognostic indicator in the metastatic group of TNBC patients (15). The higher level of CD14 in metastatic BPLER cells compared to non-metastatic HMLER cells provides additional evidence for the role of CD14 in driving an aggressiveness tumor phenotype with and an immune-cold microenvironment (Fig. 3(B)). Thus, the association of CD14 and NFkB-TGF-β could be synergistically (5,34,37).

We have previously reported distinctive restricted post-translational modifications in breast cancer which may be applicable to CD14 (38) as a potential therapeutic target.

CD14 cell membrane staining in our study, was found to be not limited to TNBC but also occurred to a lesser extent in HR+, suggesting relevance of CD14 across breast cancer subtypes, thus potentially serving as a new marker for aggressive breast cancers, and could serve as a diagnostic marker predictive of poor outcome and potentially as a therapeutic target.

## Supplementary data

**Supplementary Fig. S1.**
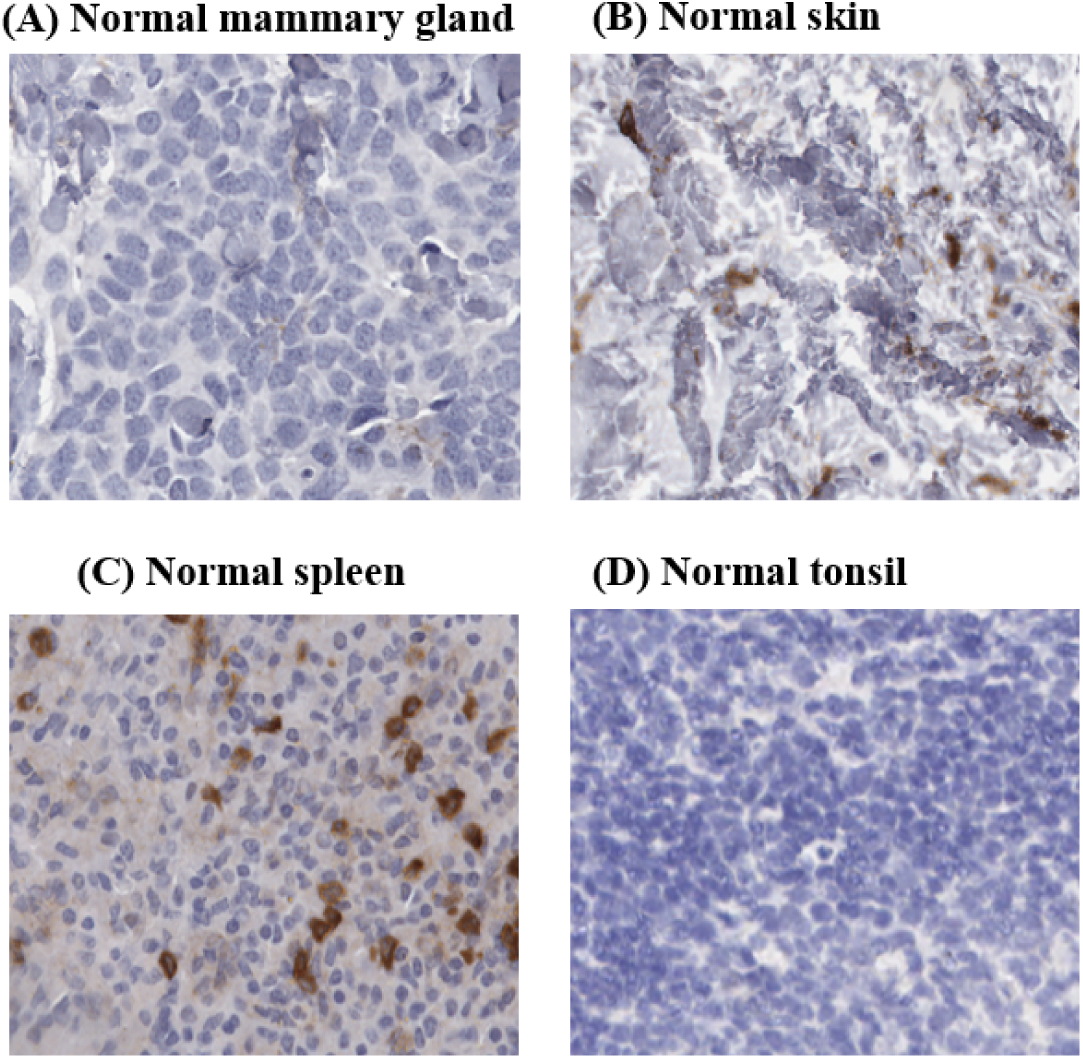
Anti-CD14 antibody IHC on normal tissues. (A) Normal mammary gland, (B) Normal skin, (C) Normal spleen and (D) Normal tonsil

**Supplementary Fig. S2.**
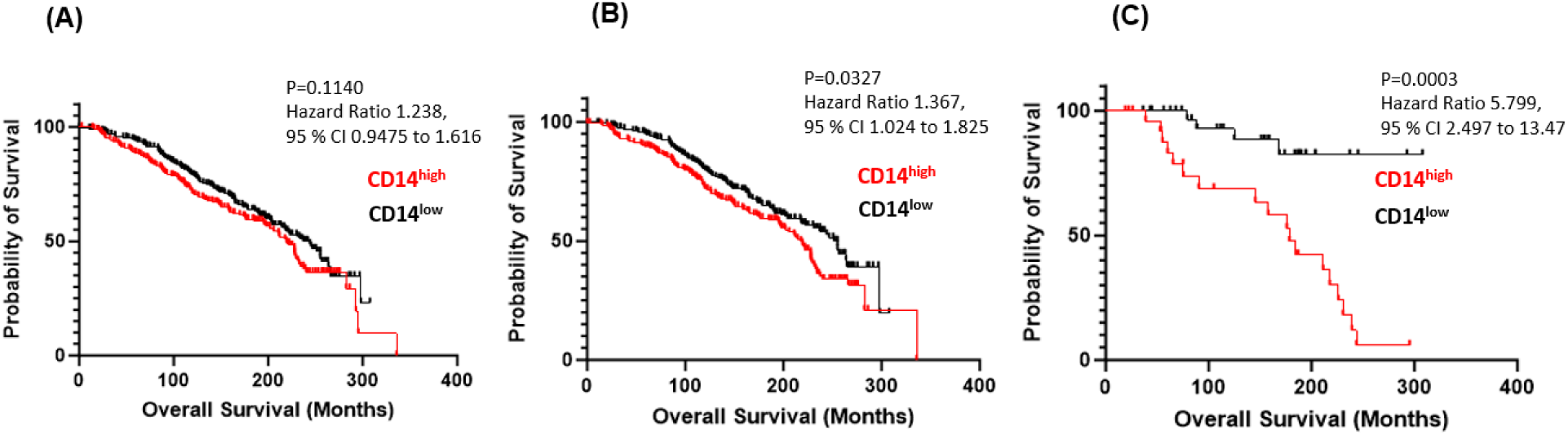
Kaplan Meier plots of gene expression in breast cancer patients (Curtis cohort) (A) HR+ and TNBC merged, stage 0 and I (n=483). (B) HR+, stage 0, and I (n=412). (C) TNBC, stage 0 and I (n=71).

## Supplementary Tables

Supplementary Table S1 Immune cell surface markers

Supplementary Table S2 Immune markers, cell surface proteome

Supplementary Table S3 Immune markers, TCE proteome

Supplementary Table S4 CD14 RNA-seq gene expression from 66 breast cancer cell lines in CCLE

Supplementary Table S5 CD14 (+) associated proteins in TCE

Supplementary Table S6 CD14 (+) associated genes in single-cell analysis

Supplementary Table S7 Clinical characteristics of the breast cancer TMA

## Acknowledgments

We thank all the collaborators in the authors list and Wei Lu, PhD IHC Laboratory Translational Molecular Pathology and the MDACC Research Histology Core Laboratory, Grant Award: CCSG P30CA016672.

## Funding

Supported through the Duncan Family Fund

## Notes

### Competing Interest Statement

The authors have declared no competing interest.

